# Grassland Ecological Compensation Policy On Pastoral Production Efficiency—Evidence From Pastoral China

**DOI:** 10.1101/2025.07.29.667359

**Authors:** Fang Ju, Wenjie Ouyang, Chengtao Zhang, Jianjun Zhang

## Abstract

Grassland Ecological Compensation Policy(GECP) in the protection of grassland ecological environment at the same time also promote the transformation and upgrading of animal husbandry, pastoral production efficiency is one of the important indicators to evaluate the sustainable development of animal husbandry, but also has a direct impact on the economic benefits of the herders. Based on field questionnaire data from 468 herding households in the pastoral areas of Inner Mongolia Autonomous Region of China, this study used two-stage semi-parametric DEA model and mediated effect model to measure herding households’ pastoral production efficiency and analyze the effect and influence mechanism of GECP on herding households’ pastoral production efficiency, respectively.The results of the study show that: (1) the overall pastoral production efficiency of herding households in Inner Mongolia is low, with a mean value of 0.43, which is mainly caused by low pure technical efficiency. (2) GECP, the number of years of education of the household head, whether or not the head of the household receives technical training, the percentage of income from herding, the number of herding machines, the evaluation of the herding socialized services and whether or not to rent pasture have a significant positive effect on the herding efficiency of herding households. (3) the realization of appropriate scale management by herding households plays a part of the mediating effect in the process of affecting herding households’ pastoral production efficiency. The realization of moderate scale operation by herders plays a partial intermediary effect in the process of GECP affecting herder’s pastoral production efficiency, and the direct effect of this policy on the pastoral production efficiency of large-scale herders is significantly larger than that of small-scale herders. Based on the above conclusions, this paper puts forward the suggestions of improving the supporting policies of ecological bonus, demonstrating and popularizing the breeding technology, and guiding the moderate scale management.

## 1. Introduction

Globally, the over-exploitation of ecosystems is still a serious problem that needs to be solved in the process of human development, and how to harmonize the promotion of ecological environmental protection and the enhancement of people’s well-being has become the focus of research by national governments ^[1]^. In this context, Payment for Ecosystem Services (PES) programs have received widespread attention as an incentive-based environmental policy tool in areas where ecologically fragile zones overlap with poverty zones^[2][3]^. The core logic is that, based on the principle of “who protects, who benefits”, the external effects of ecological services are internalized through the economic compensation mechanism, so as to achieve the dual goals of ecological protection and livelihood improvement in a synergistic manner ^[4]^. Over the past few decades, this model has evolved into a policy consensus among different countries.In Switzerland, the Ecological Compensation Area Policy (ECA) encourages farmers to set aside ecological compensation areas on their farms by granting them subsidies, which in turn promotes biodiversity restoration^[4]^. And in the United States, the Conservation Reserve Program (CRP) leaves environmentally sensitive agricultural land fallow, which reduces soil erosion, improves water quality, increases carbon sinks, and protects ecological and environmental security^[6]^.

In China, grassland is one of the most important terrestrial ecosystems, as well as an important vehicle for livestock production and a means of production on which herders depend. However, under the combined influence of extreme climatic and anthropogenic factors, grasslands have been degraded to varying degrees, seriously threatening ecological security and the livelihoods of herders^[7]^. To this end, in 2011, China began to implement a large-scale payment for ecosystem services program, the Grassland Ecological Conservation Subsidy and Incentive Mechanism Policy (hereinafter referred to as the “Grassland Ecological Compensation Policy(GECP)”), in the major grassland pastoral provinces (regions), on a five-year cycle, which has now reached its third round. This policy aims to guide herders to reduce livestock to protect the ecology and realize the sustainable development of grassland and animal husbandry by giving herders subsidies for grazing bans, incentives for grass-animal balance and subsidies for pastoral production materials^[8]^. The forbidden grazing and grass-animal balance system is the core of this policy, in which “forbidden grazing” refers to the implementation of year-round forbidden grazing and closed cultivation for seriously degraded pastures, and “grass-animal balance” refers to the government’s rationalization of the livestock carrying capacity based on the size of the pasture and the ecological situation^[9]^. After the implementation of the policy, the farming methods of herders have gradually changed from free grazing to “barn feeding or semi confinement”, and there has been a marked shift in the production methods of the pastoral industry, with a concomitant change in the pastoral production efficiency of herders. Therefore, revealing the influence mechanism of GECP on the pastoral production efficiency of herders can not only provide certain empirical support for cracking the dual problems of ecological protection in grassland pastoral areas and improvement of herder’s livelihoods, but also effectively motivate herders to improve their pastoral production efficiency and continue to promote the transformation and upgrading of grassland animal husbandry.

## 2. Literature review

Existing research on the implementation effect of GECP mainly focuses on two major aspects, on the one hand, the ecological effect assessment of the implementation of the policy, focusing on the systematic change of the key indicator of natural grassland-Normalized Vegetation Index (NDVI) ^[10][11]^. It was found that after the implementation of GECP, the comprehensive vegetation cover, fresh grass yield and unit biomass of natural grassland were improved, and the ecological carrying capacity was significantly enhanced^[13]^, while such ecological effects showed significant spatial heterogeneity, i.e., the policy had a greater impact on the quality of grassland in wet areas than in dry areas^[14]^. On the other hand, the economic effect of the policy implementation was assessed, mainly focusing on the changes in the income of herders ^[15]^. The study found that GECP not only significantly increased the total income of low-income herding households through the direct payment of ecological compensation^[16]^, but also indirectly increased the total income of herding households through the transfer of labor and expansion of herd size, in addition, the study also found that the region with a high level of per capita income of the herding households benefited more from the policy, which indicated that the policy widened the development gap within the region**^Error! Reference source not found.^**. As one of the important indicators for evaluating the sustainable development of animal husbandry, pastoral production efficiency not only characterizes the utilization of natural pasture in the process of pasture production, but also directly reflects the economic benefits of herding households. By studying the impact of the GECP on the pastoral production efficiency of herders, it is possible to reveal the impact effect of the policy on pastoral production. Some studies have found that after the implementation of the GECP, the factor productivity of professional fattening farming of meat sheep and professional fattening farming of beef cattle in pastoral areas have become a stable growth trend^[17][19]^, while some studies have also found that ecological compensation is difficult to compensate for the increase in the cost of shepherding as a result of the policy, and the longer the time of the ban on animal husbandry, the greater the decrease in the efficiency of technological progress of the herding households^[20]^.

In summary, around the implementation of GECP implementation effect of the research is mainly concentrated in the ecological and economic effects, and the impact on the pastoral production efficiency of the research results are fewer, mainly focusing on the reward policy before and after the implementation of pastoral production efficiency of the comparative study, and the conclusion of the study is still controversial, few studies using econometric modeling on the GECP on the impact of herders’ pastoral production efficiency effect and the mechanism of the effect In-depth analysis. In view of this, this paper focuses on the pastoral development effect of the GECP, focusing on answering the following questions: (1) What is the current situation of herder’s pastoral farming efficiency under the background of the implementation of GECP? (2) Does the implementation of GECP affect the pastoral production efficiency of herders, and what is the effect of the impact? (3) What role does the realization of appropriate scale management by herders play in the mechanism of GECP affecting herders’s pastoral production efficiency? (4) Is there any difference in the effect of GECP on the pastoral production efficiency of herders in different pasture sizes? Through the systematic analysis of the above questions, we hope to provide new ideas for improving the pastoral production efficiency of herders, reducing the pressure of grassland ecological protection, improving the stability of herder’s income and realizing the high-quality development of the pastoral industry.

The structure of this paper is organized as follows: Section 1 presents the introduction. Section 2 presents the Review of Literature. Section 3 presents the research materials and methods, including the theoretical framework, data sources and model construction.

Section 4 analyzes the results of the study. Section 5 is for the discussion. Section 6 is for the conclusions of the study. And section 7 is for the policy recommendations.

## 3. Materials and Methods

### 3.1. Heoretical analysis framework and research hypothesis

GECP encourages herders to protect grassland ecology through government supervision and subsidies, and at the same time drives the transformation and upgrading of grassland animal husbandry^[21]^. First of all, due to the restriction of grazing, the herder’s breeding mode gradually changed from rough free grazing to “barn feeding” or “grazing + barn feeding”, breaking the traditional grazing mode of selection of pasture and livestock management are overly dependent on the natural environment and climatic conditions of the situation. According to the theory of “induced technological innovation”, barn feeding will encourage herders to adaptively learn new breeding techniques and management methods to feed their animals scientifically, and ultimately achieve the effect of increasing the rate of slaughter and speeding up the turnover of livestock. Secondly, the policy has promoted the application and popularization of pasture machinery, especially the increase in the number of mixers and forage harvesting equipment, etc. is extremely obvious, and the increase in the level of infrastructure inputs has reduced labor inputs, providing better hardware conditions for the development of animal husbandry^[22]^. Finally, the use of pasture is also gradually transformed to the combination of natural pasture and artificial forage land, the integration of grass and livestock is obviously strengthened, the system of rest and rotational grazing improves the productivity of natural pasture, and the cultivation of high-quality pasture grasses such as shepherd’s croft, oat and alfalfa improves the self-sufficiency rate of fodder, and also provides a solid guarantee for the cold-season supplemental feeding to meet the nutritional needs of livestock and at the same time reduces the cost of fodder^[23]^. In conclusion, the policy has positively affected all aspects of herder’s production and improved the pastoral production efficiency. In view of this this paper proposes the hypothesis H1:

H1:GECP has significantly improved the pastoral production efficiency of herders. GECP promotes the reduction of herder’s herd size and the realization of moderate-scale management, which will indirectly affect herder’s pastoral production efficiency. In the past, constrained by the scale of pastoral grassland animal husbandry and the degree of specialization of herders, the scale of herder’s breeding exceeded the carrying capacity of grassland ecology, resulting in the deterioration of grassland ecology, slow increase in the income of herdsmen, and low economic efficiency of grassland animal husbandry and so on ^[24]^. One of the positive externalities of the implementation of the policy is that it prompts herders to form the production decision of moderate-scale management, which is due to the policy’s regulation of herders’ livestock carrying capacity, narrowing the gap between the existing herd size and the optimal scale^[25]^. In the process of livestock reduction, the herder optimize the allocation of breeding resources, on the one hand, the herder will improve the quality of livestock, eliminating the “inferior” livestock in the population, introducing high-quality breeds of livestock for improvement, and carrying out “selective breeding” to improve the output capacity of the unit of livestock. On the other hand, the herders optimize the breeding structure, reduce the number of livestock with poor breeding efficiency according to the market situation, and even carry out single-species “fine breeding”, and finally obtain the scale with the best operating efficiency, so as to realize the moderate scale of operation. In the moderate scale of breeding, the grassland ecosystem can reduce the adverse effects of climate change, fully release the production capacity of the natural pasture, provide sufficient forage supply for livestock, reduce the pressure of competition for food among livestock, so as to maintain good growth and reproduction capacity ^[25]^. In view of this this paper proposes hypothesis H2:(**Figure 1**)

**Figure 1.**
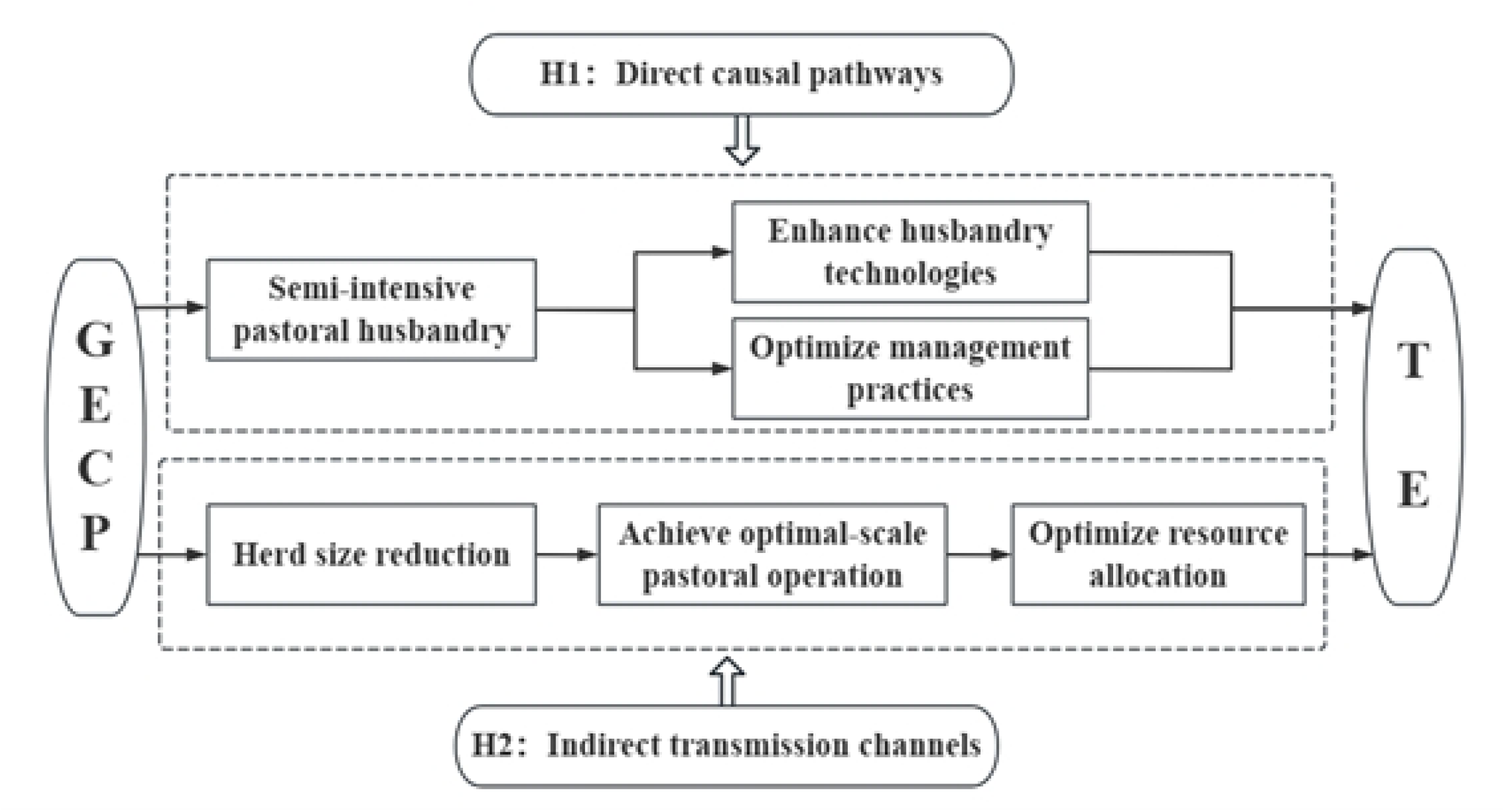
Theoretical Framework of the Impact Mechanism of GECP on Pastoral Production Efficiency

H2: Realization of appropriate scale management has a mediating effect in the path of GECP on herder’s pastoral production efficiency

### 3.2. Household surveys and data collection

The data used in this study come from field surveys conducted by the group from January to May 2024 using household surveys in the pastoral areas of Inner Mongolia, China. The Inner Mongolia Autonomous Region (IMAR) is located in the northern part of China and has a long and narrow shape, which is usually divided into 3 major parts, namely the eastern, central and western parts, based on natural geographic, climatic and economic characteristics. The survey area includes Hulunbeier City and Chifeng City in the eastern region, Xilingol League in the central region, and Ordos City in the western region, totaling four leagues (cities). The reason for the selection is that the grassland in the research area is seriously desertified, and the contradiction of “human-grass-animal” is more prominent. Firstly, stratified sampling method was used to select 1~4 banners (counties) from each league (city) according to the annual per capita net income of the county; secondly, 1~3 soums (townships) and 2~3 gacha (villages) were selected from each banner (county) according to the per capita pasture area of soum (township). And finally, 12~17 farming and herding households were selected as survey samples from each gacha (village) by simple random sampling and typical sampling methods. Finally, in each gacha (village), 12 to 17 farmers and herdsmen were selected through simple random sampling and typical sampling methods. This research in the grass-animal balance area issued a total of 510 herdsmen questionnaires and 47 village (gacha) questionnaires, after removing the questionnaires with serious missing data, a total of 468 effective herdsmen questionnaires, with an effective rate of 91.76%; 45 effective village questionnaires, with an effective rate of 95.74%. The valid sample data covers 9 herding flags (counties), 19 soums (townships) and 45 villages (gacha).

### 3.3. Model methods

#### Two-Stage Double Bootstrap DEA

The DEA model, also known as Data Envelopment Anlysis (DEA), is a nonparametric measure for evaluating the relative effectiveness of decision-making units. The method determines the nonparametric segmented front surface of the data and measures the efficiency value by constructing different decision-making units using linear programming. Characterized by the fact that there is no need to take into account the form of the production function and the magnitude of the dimension, that it is less affected by function setting errors, and that it is possible to estimate efficiencies for multi-input data and multi-output data. The models include the CRS model with constant returns to scale and the VRS model with variable returns to scale, each of which can be subdivided into two categories: input-oriented and output-oriented ^[27]^. Although most of the existing studies have used the DEA two-stage approach (DEA+Tobit) to analyze the influencing factors of pastoral production efficiency, the method may invalidate statistical inferences about the magnitude of impacts on environmental variables based on efficiency estimates because of serial correlations of complex form and unknown structure between efficiency estimates. As a result, Simar and Wilson (2007) ^[28]^ proposed a two-stage semi-parametric DEA method (Two-Stage Double Bootstrap DEA) to solve the problem, which is a hybrid method combining non-parametric efficiency analysis (DEA) and statistical inference (Bootstrap), and the core of the method is to correct the efficiency values through repeated sampling for the estimation bias and standard error of regression coefficients to more reliably analyze the influencing factors of efficiency,The approach is delineated as follows: (i) Estimate the efficiency scores of each herder using Equation (2); (ii) Estimate the truncated regression using Equation (4); (iii) calculate a set of bootstrap estimates; (iv) calculate each herders’ TE score adjusted for bias; (v) the TE scores adjusted for bias serves as the response variable in a truncated regression to estimate the factors affecting TE; (vi) conduct a series of bootstrapping to provide bootstrap estimates; and (vii) calculate new confidence intervals based on the bootstrap estimates.Therefore, this method was chosen in this paper to measure the pastoral production efficiency of the herder in the first stage and further to further analyze its influencing factors in the second stage.

In the first stage of the DEA model selection process, Coelli et al. (2005) ^[29]^ argued that a constant returns to scale CRS model is only appropriate if herders operate at an optimal scale, and that incorrect assumptions about constant returns to scale may lead to inconsistent efficiency scores and consequently to a loss of statistical efficiency. In the pastoral areas of Inner Mongolia, imperfect competition, financial, pasture area and socioeconomic constraints may prevent herder from realizing the optimal scale of herd. Therefore, the hypothesis testing procedure on returns to scale proposed by Simar and Wilson (2002) ^[30][31]^ was utilized to test the herd size profile of herder in the study area. Specifically, the bootstrap statistic 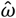 was constructed to test the original hypothesis that “H0= Overall Herd Technology Scale Reward is Constant” and the alternative hypothesis that “H1= Overall Herd Technology Scale Reward is Variable”. 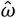 is the statistic in Simar and Wilson (2002) Equation 4.6, which has been shown to be the most powerful of all statistics.

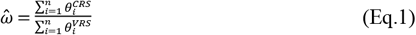

In (Eq.1), 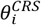 and 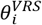 are the pastoral production efficiency values of each herder calculated by CRS model and VRS model respectively, and the test 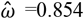, P-Value=0.03, therefore, the original hypothesis of “H0= Overall Herd Technology Scale Reward is Constant”is rejected, and VRS model is selected in the first stage to measure the herder’s pastoral production efficiency efficiency, the specific model is as follows:

Stage 1: Input-oriented VRS model.

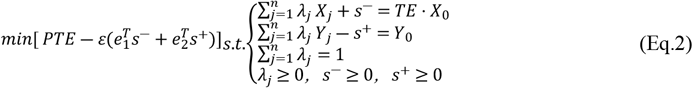

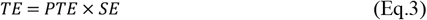

In (Eq.2) and (Eq.3), PTE stands for the pure technical efficiency of herder pastoral production. SE stands for the scale efficiency of pastoral production. And TE stands for the integrated technical efficiency of herder pastoral farming, i.e., pastoral production efficiency of herder, which is the product of the pure technical efficiency of herder pastoral production and the scale efficiency of pastoral production^[29]^. *X*_*j*_ and *Y*_*j*_ represent input and output vectors. *S*+ and *S*^−^ represent slack variables. *λ* represents a vector of weights for different herders.*j* = 1,2,3,…,*n* represents the jth herder. And *ε* represents non-Archimedean infinitesimals.

In the grassland pastoral areas of Inner Mongolia, China, after the implementation of the GECP, herders have changed their herd methods to”barn feeding” or “grazing+barn feeding” to raise livestock such as cows, sheep, and horses in the pastureland, and take the income from the sale of adult livestock, youngstock and other pastoral by-products such as wool, milk and hides will be the main source of income. At the same time, the number of livestock stocked and the number of animals slaughtered each year are relatively stable, and most of the herders will choose to carry out concentrate feed supplementation in the cold season, and some of the herders will also fatten the weaned cubs in order to make them come out of the barn in advance. Thus, drawing on the studies of Qian et al. (2019) ^[32]^ and Wang et al. (2023) ^[33]^, this paper selects herd income and year-end livestock stock as output variables, and the actual operating area of pastureland of herding households, labor inputs, and ranching operating expenses as input variables, and the specific descriptions of input and output indicators are shown in ***Table 1***.

**Table 1.** Description and descriptive statistical table of input and output indicators of livestock breeding in 2023.

| Form | Specific indicator | Indicator description | Mean | Standard deviation |
| --- | --- | --- | --- | --- |
| Input indicators | Grassland inputs | Contracted grassland area - forbidden grassland area + renting-in grassland area - renting-out grassland area (ha) | 231.41 | 217.43 |
|  | Labor inputs | Work days of own labor + work days of hired labor (work days) | 370.88 | 293.09 |
|  | Ranching operating expenses | Forage fee + medical and epidemic prevention fee + depreciation of fixed assets + energy fee (ten thousand yuan) | 14.28 | 24.43 |
| Output indicators | herd income | Herding income of herders (ten thousand yuan) | 21.72 | 23.18 |
|  | year-end livestock stock | Livestock stock at the end of the year (sheep units) | 407.30 | 390.60 |

Note: According to the Regulations on Protection of Basic Grasslands in the Inner Mongolia Autonomous Region, livestock are all converted into sheep units: 1 goat = 0.9 sheep units, 1 cow = 5 sheep units, 1 horse = 6 sheep units, 1 camel = 7 sheep units.

Stage 2: Unilateral truncated Bootstrap model.

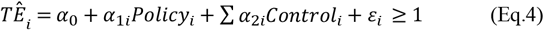

where 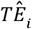 denotes the inverse of the breeding efficiency of the ith herder in Inner Mongolia pastoral area, *Policy*_*i*_ is the core explanatory variable, denotes the total amount of grassland ecological compensation received by the herders, *Control*_*i*_ is the other influencing factors, *α*_0_ is the constant term, *α*_1*i*_ is the regression coefficient, *α*_2*i*_ is the regression coefficient of the other influencing factors, and *ε*_*i*_ is the random interference term.

Explained variable: drawing on the study of Zhang et al. (2017)^[34]^, this paper takes the value of the comprehensive technical efficiency of herd of herders measured by the input-oriented VRS model as the pastoral production efficiency of herders.

Core explanatory variables: this paper selects the total amount of grassland ecological compensation in 2023 as the core explanatory variable, which is the sum of the grass-animal balance incentive and the amount of the grazing moratorium subsidy.

Control variables: drawing on the studies of Wu et al. (2021) ^[35]^and Liu et al. (2018) ^[36]^, this paper starts from the perspective of household head characteristics, the perspective of household resource endowment, the perspective of social resource endowment, and the perspective of location endowment, respectively, and selects the age of the head of the household, the number of years of education of the head of the household, whether or not the head of the household has been trained in ranching technology, the size of the household, the share of income from ranching, the number of farming machinery, and whether or not they rent into the pasture, Evaluation of socialized services in animal husbandry, whether to participate in animal husbandry insurance, and distance from the local flag (county) government were used as control variables (***Table 2***). *Mediation Effect Model*

**Table 2.**
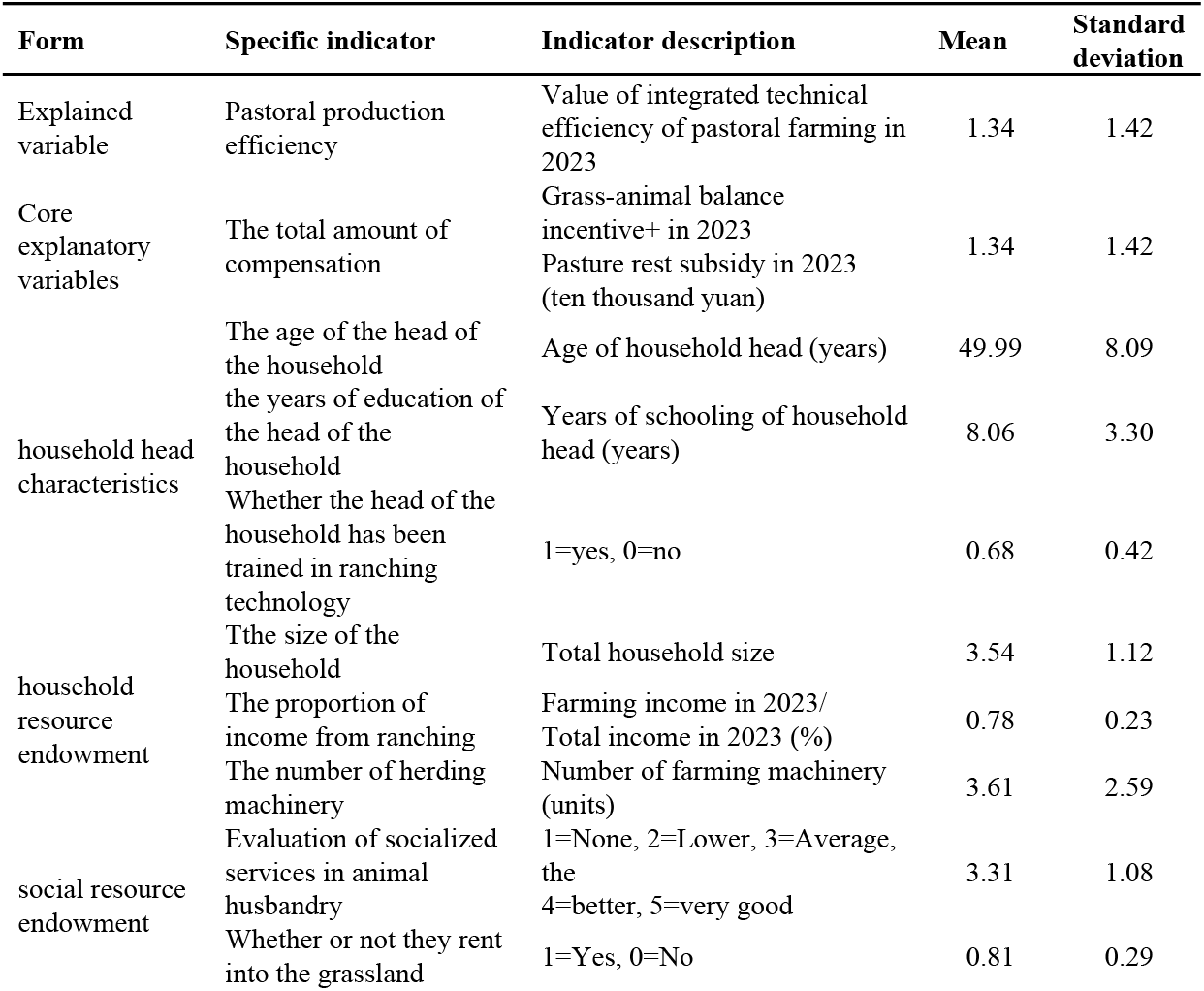
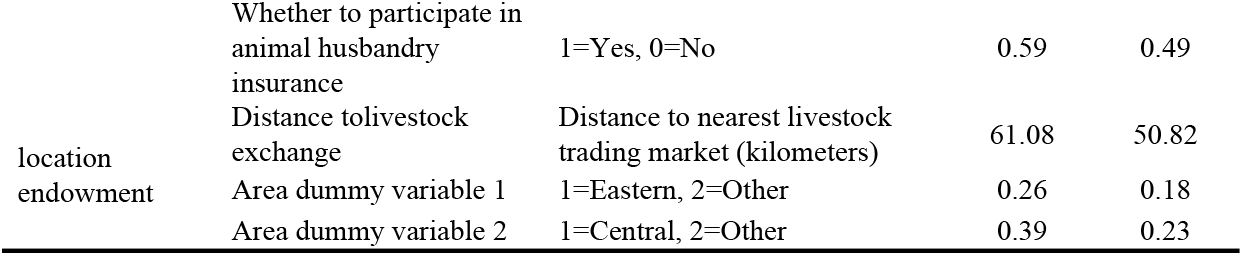
Description and descriptive statistical table of influencing factors and variables of breeding efficiency.

In order to verify whether moderate-scale operation is a mediating variable in the path of GECP affecting herders’s pastoral production efficiency, this paper adopts the mediating effect model test proposed by Wen and Ye (2014) ^[37]^to test the mechanism of the GECP’s effect on herders’s pastoral production efficiency in the form of the following model:

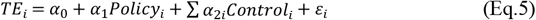

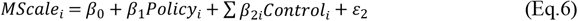

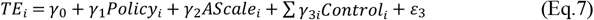

Because the pastoral production efficiency takes values in the range of [0,1], which belongs to the truncated data, (Eq.5) and (Eq.7) choose the Tobit model, and the herding households realize the appropriate scale of operation as a dichotomous variable, referring to the research of Fang et al. (2017)^[38]^, (Eq.6) chooses the Logistic model. In the above equation, *TE*_*i*_ denotes the breeding efficiency of the ith herder in Inner Mongolia pastoral area, *Policy*_*i*_ denotes the total amount of grassland ecological compensation obtained by the herder, *MScale*_*i*_ is the intermediary variable, which represents whether the herder realizes moderate scale operation, and *Control*_*i*_ is the control variable; the reasons for the selection are as follows.

The moderate scale of grassland livestock husbandry refers to the scale of breeding that realizes the win-win situation of ecological protection and economic benefits under the premise of protecting grassland ecology and combining the resource carrying capacity, economic returns and management level, and reasonably controlling the number of livestock and the mode of production^[39][40]^. At the same time, grassland as the most important resource in the process of grassland livestock production, is the basis for the allocation of other factors of production, i.e., when the breeding technology is established, the grassland resources owned by the herdsmen determine the scale of livestock production and the amount of factor inputs^[32]^. Thus, this paper refers to Zhang et al. (2012)^[41]^ study to project the moderate scale of livestock rearing based on the pasture carrying capacity operated by the herders, and since the herders’s herd size in this paper is the stock of livestock at the end of 2023, the pasture carrying capacity is referred to the “Grassland Monitoring Report of Inner Mongolia Autonomous Region in 2023” in the natural grassland cold-season livestock-carrying capacity of the various herding flags and counties, which is shown in Table 3, and the herders’s moderate The formula for calculating the herd size and the degree of deviation of herders from the moderate herd size is as follows:

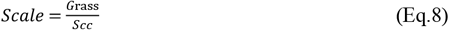

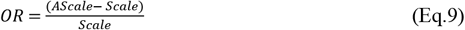

**Table 3.**
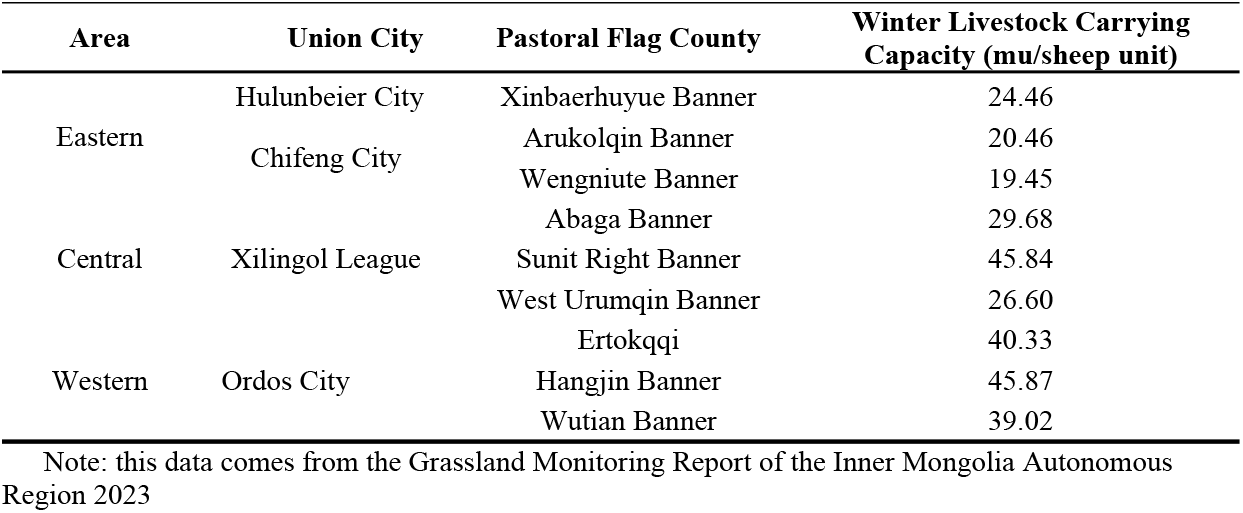
Investigate the winter carrying capacity of natural pastures in the survey area.

(Eq.8) is the formula for calculating the moderate herd size of herders, where *Scale* is the moderate herd size, *G*rass is the actual area of pasture operated by herdsmen (area of own grassland area of rented out grassland + area of rented in grassland), and *Scc*is the capacity of carrying livestock in winter (***Table 3***), and (Eq.9) is the formula for calculating the degree of herders’s deviation from the moderate herd size, where*OR* is the degree of herders’s deviation from the moderate herd size, and *AScale* is the herders’s actual herd size, measured by livestock stock at the end of 2023. In the actual management process of herders, because of the indivisibility of livestock, herders’s herd size is in a certain range of dynamic fluctuation state, this paper draws on the research results of Shi et al. (2022) ^[42]^, that herdsmen realize moderate scale operation when ―5% ≤ *OR* ≤ 5%, at this time the mediator variable *MScale* = 1, otherwise *MScale* = 0. It has been measured that, under the impetus of GECP, about 23.07% of the herders realized the moderate scale operation.

In this paper, we test the mediating effect of herders’s realization of moderate-scale operation in the mechanism of GECP on their pastoral production efficiency according to the following five steps: the first step is to test whether the core explanatory variable *Policy*_*i*_in (Eq.5) is significant. If it is significant, the argument is based on the mediating effect. Otherwise, the argument is based on the masking effect. However, no matter whether it is significant or not, the follow-up test is required. The second step is to test the core explanatory variable *Policy*_*i*_ in (Eq.6) and the mediating variable *MScale*_*i*_ in (Eq.7) in turn, if both variables are significant, then it indicates that the indirect effect is significant, and directly proceed to the fourth step. If at least one variable is not significant, then proceed to the third step. In the third step, the Bootstrap method is used to directly test 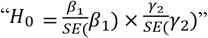, if the original hypothesis is rejected, it indicates that the indirect effect is significant, and then proceed to the fourth step. Otherwise, it indicates that the indirect effect is not significant and the analysis is stopped. The fourth step is to test whether the core explanatory variable *Policy*_*i*_ in (Eq.7) is significant, if it is not significant, it means that the direct effect is not significant, and there is only a mediating effect, i.e., there is a complete mediating effect; if it is significant, it means that the direct effect is significant, and then proceed to the fifth step of the test. In the fifth step, compare the sign of *β*_1_*γ*_2_ and *γ*_1_. If the same sign, it belongs to partial mediation effect, and mediation effect 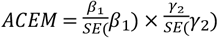. If different sign, it belongs to the masking effect, and the ratio of indirect effect to direct effect is 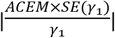.

## 4. Results

### 4.1 Analysis of Variable Returns to Scale (VRS) Model Results

In this study, the rDEA package in R language (4.4.1) was used to measure the pastoral production efficiency of herders and further analyze the mechanism of its influence. 100 interactions were conducted in the first stage of the two-stage semiparametric DEA model, and 2,000 interactions were conducted in the second stage based on Algorithm 2.

The results showed that the integrated technical efficiency, pure technical efficiency and scale efficiency of pastoral household farming biased by two-stage semiparametric DEA model all showed different degrees of decline compared with the original efficiency, and Linh and Nanseki (2015) ^[43]^in measuring the production efficiency of plantation farms in Vietnam and Tetteh et al. (2020)^[44]^in measuring the maize production efficiency of Ghanaian farm households both showed a a similar situation, as shown in ***Table 4*** and ***Figure 2***. The combined technical efficiency is the product of pure technical efficiency and scale efficiency, reflecting the overall efficiency of a pastoralist household considering both technical management and herd size at the current level of technology. The mean value of the bias-corrected integrated technical efficiency value is 0.43, and only 8% of the herders are in the high efficiency range, which indicates that the herders’ inputs need to be reduced by about 57% in order to raise the integrated technical efficiency value to 1. Meanwhile, the lower and upper mean values of the bias-corrected integrated technical efficiency value are 0.41 and 0.45, respectively, in the 95% confidence interval, which indicates that an average herder can save about 55%-59% of the total inputs by improving the integrated technical efficiency. The pure technical efficiency of the herders’s farming is the farming efficiency of the herders under the existing technical level and management ability after excluding the influence of scale factor; after calculation, the mean value of the pure technical efficiency value corrected by the herders is 0.49, and only 2% of the herders are in the high-efficiency interval, which indicates that the herders can still improve 51% of the pure technical efficiency by adopting the advanced farming technology and management. Meanwhile, the value of the pure technical efficiency value corrected by the deviation is 0.49 within the 95% confidence interval, and its lower limit and upper limit mean value are 0.42 and 0.52 respectively. The herd size efficiency of herders reflects the gap between the existing herd size of herders and their theoretical optimal herd size under the existing resource endowment and management ability. By calculation, the average value of the size efficiency of herders after corrective bias is 0.86, and about 78% of herders are in the high-efficiency interval, while the size efficiency value after bias is within the 95% confidence interval, and the lower and upper limits of the average value are 0.81 and 0.94, which indicates that GECP plays an important role in making herders get rid of the mismatch between the past herd size and grassland productivity, and realize the “grass for livestock” principle.

**Table 4.**
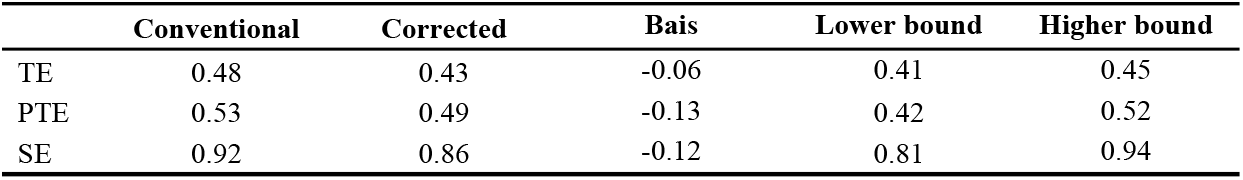
Efficiency measurement results table.

**Figure 2.**
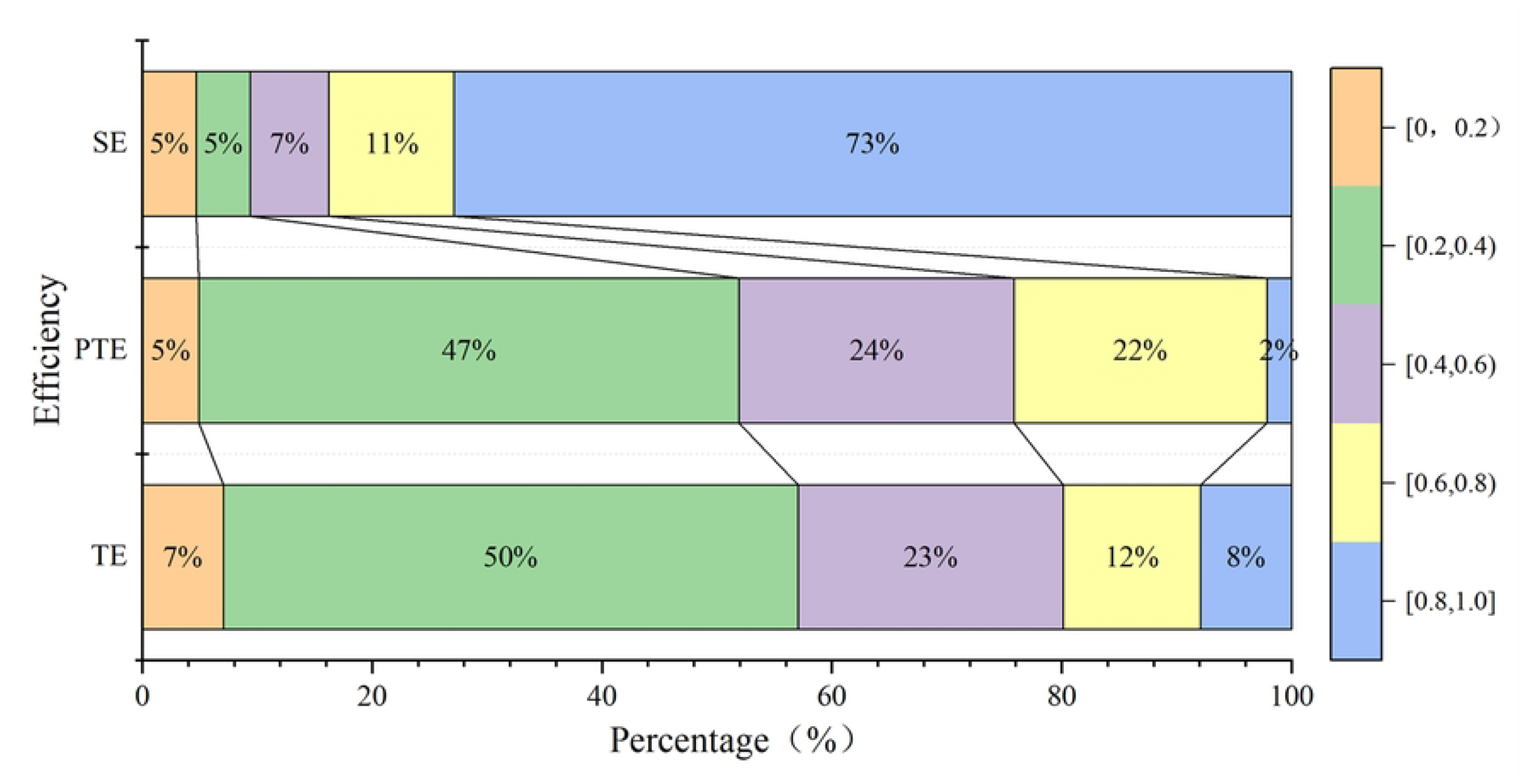
Distribution of Pastoral Production Efficiency Scores

### 4.2. Single-Truncated Bootstrap Regression Analysis

Before regressing the unilateral truncated Bootstrap model, the variables were first tested for multicollinearity using the Variance Inflation Factor Test, which showed that the largest variance inflation factor was 1.08 and the average variance inflation factor was 1.04, which is much less than 10, so there is no serious multicollinearity problem. The core explanatory variable, the total amount of grassland ecological compensation, has a significant positive effect on the pastoral production efficiency of herders in Inner Mongolia pastoral areas at the 1% level (***Table 5***), and hypothesis H1 is verified. This is consistent with the findings of Ma et al. (2021) ^[25]^ and Wang et al. (2021)^[45]^, who concluded that GECP will, on the one hand, GECP will prompt herdsmen to change their traditional farming methods from free-ranging to “barn feeding or semi confinement”, and herders will take the initiative to improve their own farming technology and management ability to adapt to the new farming methods, thus improving the pastoral production efficiency. On the other hand, GECP will constrain the scale of herders’s farming through the stipulation of the amount of livestock carrying capacity, and overloaded herders will optimize the structure of the livestock population during the process of livestock reduction, eliminate old, weak and sick animals, and actively introducing good livestock breeds for improvement, so as to enhance the production capacity of each unit of livestock, and thus improve the pastoral production efficiency. Moreover, after obtaining the bonus amount of grassland ecological bonus, the herders will improve the standardized and mechanized level of breeding by improving the conditions of the facilities of the shed and purchasing additional breeding machinery, etc, and ultimately improve the pastoral production efficiency.

**Table 5.** Regression result table of unilateral stage Bootstrap model.

| Variable | Regression | 90% confidence interval | 95% confidence interval | 99% confidence interval |
| --- | --- | --- | --- | --- |
| Bonus amount | -0.2204*** | [-0.3604,-0.0743] | [-0.4674,-0.0860] | [-0.4382,-0.0562] |
| Age of household | -0.0010 | [-0.2999,0.02570] | [-0.0312,0.0305] | [-0.0382,0.0418] |
| Years of education of household | -0.1139* | [-0.1831,-0.0306] | [-0.1859,0.0157] | [-0.2210,0.0434] |
| Whether household head receives technical training | -0.1448** | [-0.3535,-0.2045] | [-0.3853,-0.0705] | [-0.5111,0.2488] |
| Size of family | -0.2558 | [-0.4148,0.0556] | [-0.4356,0.1486] | [-0.5080,0.2028] |
| Percentage of income from herding | -3.7967*** | [-4.6801,-2.8148] | [-4.9177,-2.5394] | [-0.5472,-2.3379] |
| Number of herding machinery | -0.1144** | [-0.2195,-0.0122] | [-0.2535,-0.0302] | [-0.2360,0.0520] |
| Evaluation of herding socialization services | -0.5193** | [-0.7005,-0.3508] | [-0.7766,-0.2851] | [-0.8194,0.0903] |
| Whether pasture is leased | -1.0919** | [-1.7853,-0.4566] | [-1.7866,-0.4398] | [-2.0367,0.2341] |
| Whether pasture is insured | -0.2815 | [-0.7773,0.2332] | [-0.7893,0.2581] | [-0.7913,0.5032] |
| Distance to livestock market | 0.0021 | [-0.0014,0.0060] | [-0.0025,0.0075] | [-0.0029,0.5032] |
| Regional dummy variable1 | -0.1286 | [-0.3148,0.0556] | [-0.3546,0.1538] | [-0.4148,0.1918] |
| Regional dummy variable1 | 0.2568 | [-0.0148,0.3532] | [-0.1325,0.3968] | [-0.1958,0.4015] |
| Constant term | 9.2715 | [7.0269,11.1200] | [6.8858,11.2289] | [6.5883,12.2265] |

Note: A variable is significant at a certain confidence level if its regression coefficient does not include zero in the confidence interval at that level; the dependent variable for this regression is the inverse of the farming efficiency of the herdsmen.

Among the control variables, in terms of the characteristics of the head of household, the number of years of education of the head of household has a significant positive effect on the farming efficiency of the herding households at the 10% level, indicating that the higher the level of education of the herding households, the stronger their ability to learn and acquire information, and the easier it is to master the advanced farming technology, so as to rapidly improve the pastoral production efficiency. Whether or not the head of household receives technical training has a significant positive effect on the pastoral production efficiency of the herding households at the 5% level. Nowadays, most of the herding households in the pasture area have the farming method of “grazing + barn feeding”, so mastering the corresponding foddering farming skills is crucial for herding households to improve their pastoral production efficiency. From the point of view of household resource endowment, the proportion of pastoral income has a significant positive impact on the pastoral production efficiency of herding households at the 1% level, which may be due to the fact that the larger the share of farming income in the household income of herding households, the higher the herding households’ dependence on farming and the higher the degree of importance they attach to farming, which makes it more likely that they can improve their pastoral production efficiency by adopting advanced farming technology and increasing farming investment. The number of pastoral machinery has a significant positive impact on the pastoral production efficiency of herding households at the 1% level. The use of mechanization is an important indicator of the level of modernization of the pastoral industry. In the pastoral area each herding household raises an average of 400 sheep units, and the use of pastoral machinery, such as mixers, shears, and haying machines, can greatly reduce labor inputs and improve the pastoral production efficiency. From the point of view of social resource endowment, the evaluation of pastoral socialization service has a significant positive impact on the pastoral production efficiency of herding households at the 5% level, that is, the higher the evaluation of local pastoral socialization service by herding households means the higher the degree of local pastoral socialization service, and the support provided by the pastoral socialization service institutions in terms of disease prevention and control guidance, market information and sales channels is relatively comprehensive, which is conducive to reducing the breeding risk and increasing the pastoral production efficiency of herding households. Whether or not the pasture is rented has a significant positive effect on pastoralists’ farming efficiency at the 5% level, on the one hand, GECP directly increases the income of herdsmen through subsidies, causing some herdsmen to quit farming and move to the city. On the other hand, herders who rent in pastureland can realize large-scale farming, which not only reduces livestock breeding and marketing costs, but also facilitates herder’s introduction of more advanced pasture machinery, breeding techniques and management tools, and ultimately improves the pastoral production efficiency.

### 4.3. Analysis Robustness Checks

In order to further test the robustness of the results of the two-stage semiparametric DEA model, this paper refers to the study of Long et al. (2020)^[46]^, and applies the traditional two-stage model (DEA+Tobit) to carry out the test, in which the dependent variable is the value of the original pastoral production efficiency measured by the DEA model, and the regression coefficients of the explanatory variables are positive, which indicates that the variable has a positive influence on the pastoral production efficiency of the herder. The regression results of the Tobit model are shown in ***Table 6***, and it can be seen that the direction of the influence of the explanatory variables on the dependent variable of this model and the two-stage semiparametric DEA model are consistent, and only the significant level of the two variables of the evaluation of pasture socialisation services and whether or not to rent into the pasture are slightly different, so the conclusions in **Table 5** are more reliable.

**Table 6.** Tobit model regression results table.

| variable | coefficient of regression | Standard error | t value | p value |
| --- | --- | --- | --- | --- |
| Total amount of supplementary prize | 0.0188*** | 0.0070 | 2.639 | 0.0083 |
| Age of head of household | 0.0006 | 0.0012 | 0.539 | 0.5898 |
| Number of years of education of the head of household | 0.0052* | 0.0029 | 1.785 | 0.0743 |
| Does the householder receive technical training? | 0.0279** | 0.0118 | 2.360 | 0.0183 |
| Family population size | 0.0078 | 0.0086 | 0.902 | 0.3671 |
| Proportion of animal husbandry income | 0.1982*** | 0.0434 | 4.546 | 0.0005 |
| Quantity of animal husbandry machinery | 0.0088** | 0.0038 | 2.359 | 0.0183 |
| Evaluation of socialization service of animal husbandry | 0.0247*** | 0.0091 | 2.679 | 0.0070 |
| Whether to rent into grassland | 0.0937*** | 0.0248 | 3.779 | 0.0002 |
| Whether to take animal husbandry insurance | 0.0185 | 0.0199 | 0.927 | 0.3541 |
| Distance from livestock trading market | -0.0001 | 0.0002 | -0.250 | 0.8024 |
| Region virtual variable 1 | 0.0433 | 0.1221 | 1.951 | 0.1510 |
| Regional virtual variable 2 | -0.0437 | 0.0201 | -0.211 | 0.8327 |
| constant term | -0.0347 | 0.0934 | -0.371 | 0.7107 |
| LR chi2(11) |  | 71.96 |  |  |
| Prob>chi2 |  | 0.0000 |  |  |

Note: \* means 10% significance level, \* \* means 5% significance level and \* \* \* means 1% significance level, the same below.

### 4.4. Validation of Mediation Effect Models

In this paper, according to the Mediating effect model test method, the results show that the overall fitting effect of the Mediating effect model is better (***Table 7***). the model results of formula (5) show that the total amount of GECP has a significant positive impact on the breeding efficiency of the herder at the 1% level, i.e., the core explanatory variable of the total amount of grassland ecological subsidy is significant, which indicates that there is a mediating effect. The results of the model in Eq. (6) show that the total amount of grassland ecological bonus has a significant positive effect on whether the herder realise moderate scale management at the 5% level, i.e. the mediating variable in Eq. (6) is significant, i.e. the herder’s ‘whether to realise moderate scale management’ is significant, and at the same time, the results of the model in Eq. (7) show that the herders’s realisation of moderate scale management has a significant positive effect on their pastoral production efficiency at the 10% level, which means that there are mediating effects. At the same time, the results of the model in Eq. (7) show that herders’s realisation of moderate-scale management has a significant positive effect on their farming efficiency at the 10% level, i.e., the mediator variable in Eq. (7) is significant, which indicates that the indirect effect is significant. After adding the mediator variable herders’s realisation of moderate-scale management in the model of Eq. (7), the core explanatory variable total grassland ecological subsidy is also significant, i.e., the core explanatory variable total grassland ecological subsidy in Eq. (7) is significant, which indicates that the direct effect is significant. In the above regression results, both and are positive, that is, the indirect effect of herders’s realisation of moderate-scale management on their pastoral production efficiency is in the same direction as the direct effect, indicating that there is a part of the mediating effect, and the mediating effect accounts for 18.42% of the total effect. Therefore, GECP has the mechanism of ‘GECP - herder realise appropriate scale management - improve pastoral production efficiency’, and hypothesis H2 is verified. In the past, due to the constraints of the scale of grassland animal husbandry and the degree of specialisation of herders, herd size exceeded the capacity of grassland ecology, which led to the deterioration of grassland ecology, slow income increase of herders, and low economic efficiency of grassland animal husbandry, and other problems. GECP through the issuance of ecological bonus and strengthen the supervision of the way to guide the herders to reduce the herd size, so that herd size and grassland production capacity to match, to avoid the phenomenon of scale of the uneconomical, in the process of herders will be based on their own grassland carrying capacity to re-adjust the input of resources, optimise the allocation of resources, to achieve the appropriate scale of operation, the formation of a pasture restoration The positive cycle of ‘pasture restoration - pasture quality improvement - livestock efficiency increase’ was formed. Li et al. (2021)^[47]^ also found that when the actual livestock loading rate of herders was reduced to the optimal loading rate, although the total expenditure and total income of herders decreased, their net income and net income per unit of livestock increased. Thus, pastoralists improved their pastoral production efficiency by achieving appropriate scale of operation.

**Table 7.** Test Results Table of the Mediation Effect Model.

| model | Tobit model | Logistic model | Tobit model |
| --- | --- | --- | --- |

| variable | TE | Mscale | TE |
| --- | --- | --- | --- |
| Policy | 0.0380***<br>(0.006) | 0.3641**<br>(0.074) | 0.0310***<br>(0.007) |
| Mscale | -- | -- | 0.0418*<br>(0.023) |
| Control variable | Controlled | Controlled | Controlled |
| Pseudo R<sup>2</sup>/ R<sup>2</sup> | 0.168 | 0.151 | 0.173 |

### 4.5. Validation of Mediation Effect Models

Grassland resources, as the most important production factors in the farming activities of herders, are not only the approved criteria for the release of ecological compensation, but also the core variable for calculating the theoretical livestock carrying capacity. According to the theory of resource endowment, in the context of the implementation of GECP, pasture resources directly determine the adjustment space of the production behaviour of the herders and the potential of the improvement of pastoral production efficiency. Therefore, in order to further analyse the heterogeneous impact of GECP on the pastoral production efficiency of herding households, this paper divides the total sample into a large herding household sample (the actual area of pasture operated by herding households is larger than the average) and a small herding household sample (the actual area of pasture operated by herding households is smaller than the average) according to the actual area of pasture operated by the herding households in 2023, and establishes the Mediating effect model to analyse the two samples separately, and the obtained results are As shown in ***Table 8***.

**Table 8.**
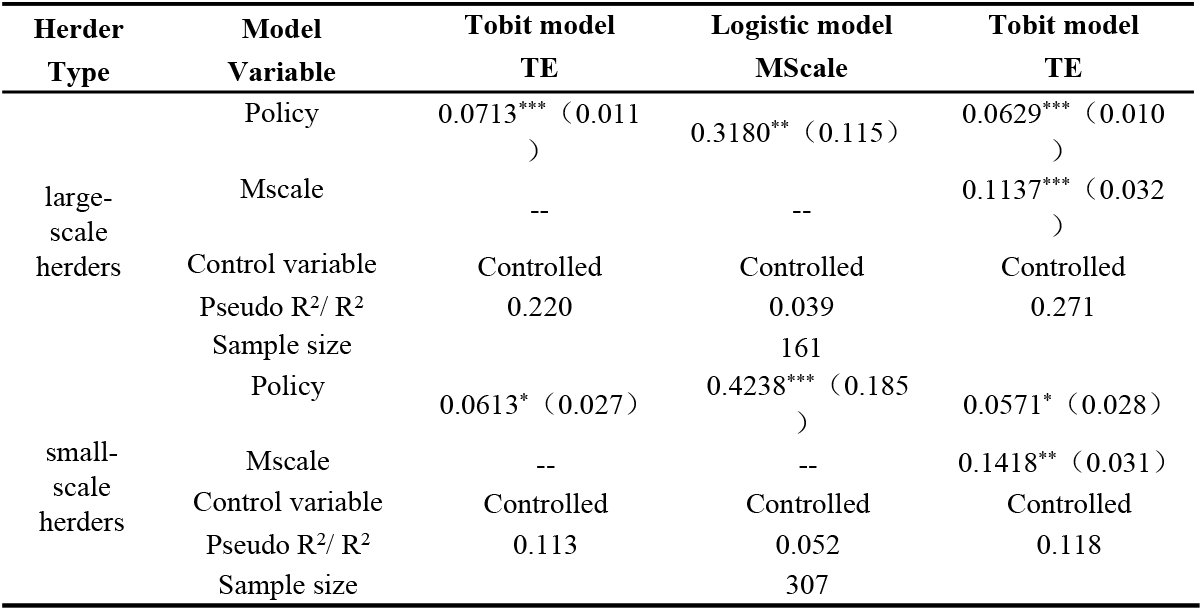
The test result table of heterogeneous herders ‘mediating effect model.

It was found that the impact of GECP on the pastoral production efficiency of herding households was still dominated by the direct effect, both in the sample of large herding households and in the sample of small herding households, and that the direction of the mediation transmission of the moderate-scale operation was in the same direction of the direct effect, which was manifested as a partially mediated effect, which was largely consistent with the results of the overall sample. Further analysis found that the indirect effect of GECP on the pastoral production efficiency of small-scale herders (22.34%) is significantly larger than that of large-scale herders (11.78%), which may be due to the fact that small-scale herders used to be the main body of overloaded grassland pasture areas, and the resource allocation of this part of herders is in an imbalanced state. While the GECP promotes small-scale herders to reduce the scale of their breeding by means of supervision and subsidies, and guides them to control the herd size in the carrying capacity of the pasture. GECP promotes small-scale herding households to reduce their herd size through supervision and subsidy, guides them to control their herd size within the carrying range of pasture, improves the matching degree between their pasture resources and herd size, solves the previous vicious circle of ‘pasture degradation - overgrazing’, and indirectly improves the pastoral production efficiency by realising the appropriate scale of operation. Comparatively speaking, large-scale herders have richer pasture resources, and these herding households are better than small-scale herders in terms of both breeding scale and the total amount of subsidies received, so they will increase investment in breeding, such as sheltered sheds and breeding machinery, and thus improve breeding technology and management level, and the direct effect of GECP on pastoral production efficiency is even greater. This pattern of differentiation suggests that policy design needs to take into account the characteristics of the group, and that small-scale herders should work together to strengthen the supervision of grass-animal balance and support alternative livelihoods, while large-scale herders need to guide the flow of compensatory funds to technological innovation, so as to ultimately realise the scale-matched development path of ecological protection and the transformation of the pastoral industry.

## 5. Discussion

Based on 468 field research questionnaires in pastoral areas of Inner Mongolia, this paper empirically analyses the effect of GECP on the pastoral production efficiency of herders and the influence mechanism on the pastoral production efficiency of herders, and further clarifies the mechanism of herders’s realisation of the role of the appropriate scale of operation in the relationship between the two. The results found that in the pastoral areas of Inner Mongolia, China, although GECP has improved the pastoral production efficiency of herders, but the policy effect is more limited, the current breeding efficiency of herders in this pastoral area as a whole is still in the lower efficiency range, mainly due to the lower pure technical efficiency, the pastoral production efficiency is still a large room for improvement, which is consistent with the conclusions of some of the previous studies^[34]**Error! Reference source not found**.^. The incentive effect of the implementation of GECP on the development of the pasture industry is due to the standardisation of the use of pasture, guiding herders to change to the modernised farming method of “grazing + barn feeding”, and at the same time, the ecological compensation issued compensates for the production costs incurred during the early stage of the transformation of herders’s farming, so that the herdsmen will adapt to the new way of farming and learn advanced farming techniques and management methods, which directly improves their farming efficiency. Under the new farming method, the herders will adapt to learn advanced farming techniques and management methods, which will directly improve the herder’s pastoral production efficiency. However, it cannot be ignored that the pastoral areas of Inner Mongolia are sparsely populated, and still face the same “aging population” dilemma as China’s agricultural areas**^Error! Reference source not found.^**, resulting in the slow diffusion and adoption of farming management techniques, which seriously restricts the economic development of pastoral areas.

Through the mechanism test, this paper found that the herders realised the moderate scale operation in GECP affects the herders’s pastoral production efficiency process to play a part of the mediating effect, and the policy on the small herders’s pastoral production efficiency of the indirect effect is significantly larger than that of the large herders. In the past, the grass-animal relationship in Inner Mongolia’s pastoral areas was not always in a state of imbalance, and its evolution can be divided into four major stages due to the influence of climate and policy: Stage 1: the pasture surplus stage of “Grass > Livestock” before 1965. Stage 2: the “Grass ≈ Livestock” stage from 1966-1985Second stage: 1966-1985 “Grass ≈ Livestock” equilibrium stage. Stage 3: 1986-2005 “Livestock > Grass” imbalance stage. Stage 4: after 2006 “Livestock > Grass” imbalance recovery stage**^Error! Reference source not found.^**. In pastoral areas, the dynamic balance of grass and livestock is the basis for high-efficiency farming. With the increasing ecological compensation and supervision, the imbalance of “livestock>grass” is being recovered at a faster pace, and herders are gradually realising appropriate scale operation and optimising resource allocation, which indirectly improves their pastoral production efficiency. At the same time, compared with large-scale herding households, small-scale herding households are more likely to fall into the serious imbalance of “livestock > grass”^[50]^, under the influence of GECP, herding households gradually transfer part of the family labour force to other industries and reduce the scale of breeding, so as to achieve the optimal allocation of breeding resources with the core of the pasture, so that the policy has a positive impact on small-scaleherding households’ pastoral production efficiency is more prominent in the indirect impact of the policy.

In order to achieve the policy objective of synergistic development of ecology, production and life in grassland pastoral areas and the dynamic balance of “people, grass and animals”, the key lies in improving the pastoral production efficiency, accelerating the transformation and upgrading of grassland animal husbandry, and thus improving the economic benefits of herders.Only when herders achieve sustainable livelihoods will it be easier to motivate them to protect grassland ecology. Research data show that 67.30% of herding households are willing to take the initiative to contribute to the protection of grassland ecology. When the grassland ecology is gradually restored, productivity will continue to increase, the development of the pastoral industry will accelerate, and a virtuous cycle of “pastoral development, increased income for herders, and ecological restoration” will ultimately be realised under the impetus of the subsidy and reward policy (*Fig 3*).How to ensure that this cycle operates more efficiently is the direction for optimising the fourth round of grassland ecological bonus policies. It is undeniable that this study is confined by the difficulty of research and only selected cross-sectional data for the study, which is both insufficient and a direction for future research in co-operation with related scholars.

**Figure 3.**
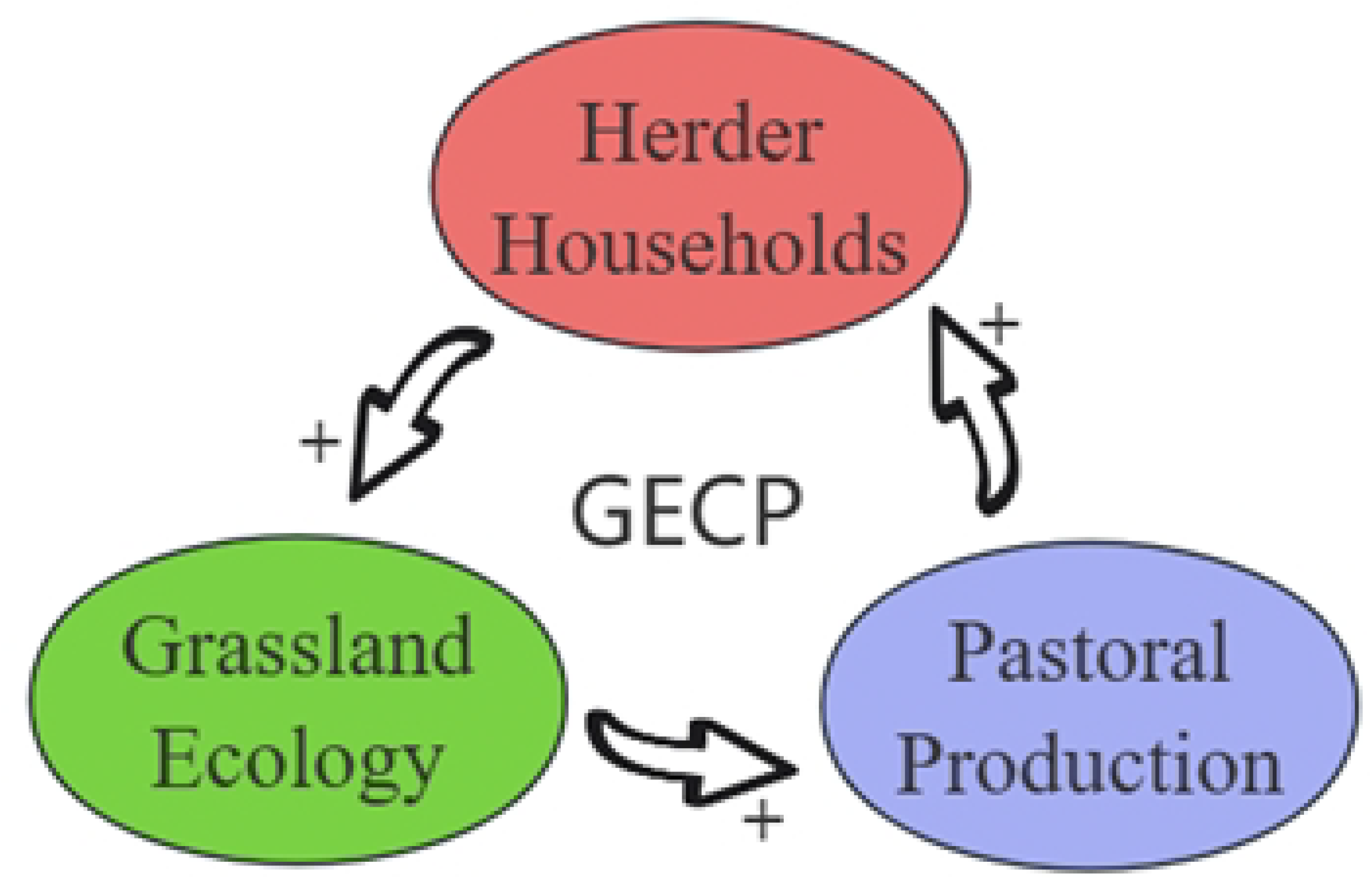
Multidimensional Effects of GECP Implementation

## 6. Conslusion

This paper is based on 468 questionnaire data from herders in the pastoral areas of Hulunbeier City, Inner Mongolia, Xilingol League, Chifeng City and Ordos City obtained from field research during the implementation period of the third round of GECP, and firstly, the two-stage semiparametric DEA model was used to measure the integrated technical efficiency, pure technical efficiency and pastoral production efficiency of the herders’s farming respectively, and then the integrated technical efficiency was used as the representative of the herder’s pastoral production efficiency, and then the integrated technical efficiency was used to represent the herders’s pastoral production efficiency.Then the comprehensive technical efficiency values were used to represent the pastoral production efficiency of the herders, and then the impact effect and influence mechanism of GECP on the pastoral production efficiency of the herders were investigated separately, and finally whether the moderate scale operation was realised or not was taken as the mediator variable, and the mediator effect model was used to verify the influence mechanism of GECP on the pastoral production efficiency of the herders and to obtain the following conclusion:

1. The pastoral production efficiency level of herders in the pastoral areas of Inner Mongolia, China, is on the low side as a whole, with a mean value of 0.43, and there is an efficiency loss of 57%, which has a large room for improvement; the mean value of the pure technical efficiency of farming is 0.49, and the mean value of the scale efficiency is 0.86; the lower level of the pastoral production efficiency is mainly due to the lower level of the pure technical efficiency of farming.
2. GECP significantly promotes the improvement of herders’s pastoral production efficiency, and among the control variables, the number of years of education of the head of household, whether the head of household receives technical training, the proportion of income from herding, the number of herding machinery, the evaluation of herding socialisation services and whether the pasture is rented have a significant positive effect on the herders’s pastoral production efficiency.
3. Herders’s realisation of an appropriate scale of management of herd size plays a part in the mediating effect in the mechanism of GECP’s influence on herder’s pastoral production efficiency, with the mediating effect accounting for 18.42%.In addition, the results of the heterogeneity analysis showed that the direct effect of GECP on the pastoral production efficiency of large-scale herding households was significantly greater than that of small-scale herding households.

## 7. Recommendations

- Improve the supporting policies of ecological subsidies and awards, and promote the transformation and upgrading of the animal husbandry industry.In the pastoral areas of Inner Mongolia, China, the “grazing + barn feeding” farming method of animal husbandry has higher requirements for supporting facilities, therefore, governments at all levels should strengthen the basic supporting security, and further improve the construction of sheds, artificial grass planting, the purchase of breeding machinery and other supporting policies related to the complementary policy support efforts,actively breaking down the financial and technical constraints faced by herdsmen at the initial stage of the transformation of their farming methods, enhancing herders’s controllability of barn feeding farming, effectively improving herders’s farming conditions, and smoothly promoting the transformation and upgrading of the animal husbandry industry.
- Demonstrate and promote breeding techniques to enhance the effectiveness of pastoral production. Guiding herders to adopt advanced breeding technology and scientific management is the key to improving pastoral production efficiency, so the level of socialized services in the pastoral industry should be continuously improved, and animal husbandry and veterinary technical service centers around the world can make use of herders’s free time to carry out training in pastoral technology, teaching and explaining the knowledge of forage proportioning, seed improvement and disease prevention and control, etc., so as to make herder’s comprehensive quality be able to adapt to the requirements of the development of the modern animal husbandry industry.
- Implementing grass-based animal husbandry and guiding moderate-scale management. Grass-animal imbalance is one of the important factors leading to the deterioration of grassland ecology and the slow development of animal husbandry, therefore, the relevant departments should approve the differentiated moderate scale of operation standards in accordance with the demands of the herders and the grassland ecological situation in sub-regions, publicize the concept of sustainable development of animal husbandry to the herders and encourage the herders to consolidate the breeding resources represented by the grassland through setting up the cooperatives or the family pasture, so as to enhance the efficiency of resource allocation, ultimately realizing improved pastoral production efficiency.

## Acknowledgements

This research was funded by the National Natural Science Foundation of China Project “Study on the Influence Mechanism of Grassland Ecological Compensation and Reward Policy on Part-time behavior of Grazing Herdsmen——Taking Inner Mongolia as an Example” (72363025)

